# Striking Antibody Evasion Manifested by the Omicron Variant of SARS-CoV-2

**DOI:** 10.1101/2021.12.14.472719

**Authors:** Lihong Liu, Sho Iketani, Yicheng Guo, Jasper F-W. Chan, Maple Wang, Liyuan Liu, Yang Luo, Hin Chu, Yiming Huang, Manoj S. Nair, Jian Yu, Kenn K-H. Chik, Terrence T-T. Yuen, Chaemin Yoon, Kelvin K-W. To, Honglin Chen, Michael T. Yin, Magdalena E. Sobieszczyk, Yaoxing Huang, Harris H. Wang, Zizhang Sheng, Kwok-Yung Yuen, David D. Ho

## Abstract

The Omicron (B.1.1.529) variant of SARS-CoV-2 (severe acute respiratory syndrome coronavirus 2) was only recently detected in southern Africa, but its subsequent spread has been extensive, both regionally and globally^1^. It is expected to become dominant in the coming weeks^2^, probably due to enhanced transmissibility. A striking feature of this variant is the large number of spike mutations^3^ that pose a threat to the efficacy of current COVID-19 (coronavirus disease 2019) vaccines and antibody therapies^4^. This concern is amplified by the findings from our study. We found B.1.1.529 to be markedly resistant to neutralization by serum not only from convalescent patients, but also from individuals vaccinated with one of the four widely used COVID-19 vaccines. Even serum from persons vaccinated and boosted with mRNA-based vaccines exhibited substantially diminished neutralizing activity against B.1.1.529. By evaluating a panel of monoclonal antibodies to all known epitope clusters on the spike protein, we noted that the activity of 17 of the 19 antibodies tested were either abolished or impaired, including ones currently authorized or approved for use in patients. In addition, we also identified four new spike mutations (S371L, N440K, G446S, and Q493R) that confer greater antibody resistance to B.1.1.529. The Omicron variant presents a serious threat to many existing COVID-19 vaccines and therapies, compelling the development of new interventions that anticipate the evolutionary trajectory of SARS-CoV-2.

## Main text

The COVID-19 (coronavirus disease 2019) pandemic rages on, as the causative agent, SARS-CoV-2 (severe acute respiratory syndrome coronavirus 2), continues to evolve. Many diverse viral variants have emerged (**Fig. 1a**), each characterized by mutations in the spike protein that raise concerns of both antibody evasion and enhanced transmission. The Beta (B.1.351) variant was found to be most refractory to antibody neutralization^4^ and thus compromised the efficacy of vaccines^5-7^ and therapeutic antibodies. The Alpha (B.1.1.7) variant became dominant globally in early 2021 due to an edge in transmission^8^ only to be replaced by the Delta (B.1.617.2) variant, which exhibited even greater propensity to spread coupled with a moderate level of antibody resistance^9^. Then came the Omicron (B.1.1.529) variant, first detected in southern Africa in November 2021^3,10,11^ (**Fig. 1a**). It has since spread rapidly in the region, as well as to over 60 countries, gaining traction even where the Delta variant is prevalent. The short doubling time (2-3 days) of Omicron cases suggests it could become dominant soon^2^. Moreover, its spike protein contains an alarming number of >30 mutations (**Fig. 1b and Extended Data Fig. 1**), including at least 15 in the receptor-binding domain (RBD), the principal target for neutralizing antibodies. These extensive spike mutations raise the specter that current vaccines and therapeutic antibodies would be greatly compromised. This concern is amplified by the findings we now report.

**Fig. 1.**
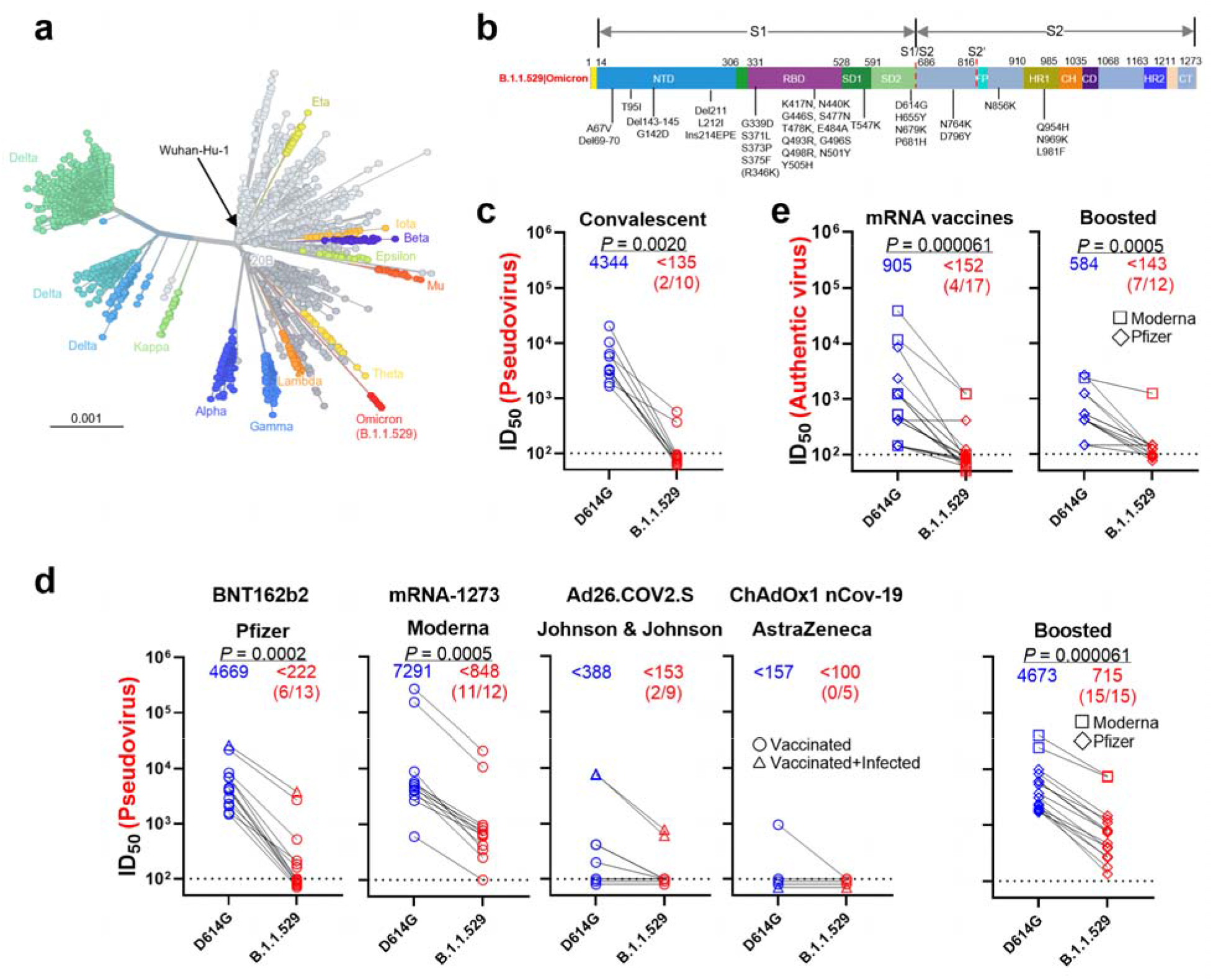
Resistance of B.1.1.529 to neutralization by sera. **a**, Unrooted phylogenetic tree of B.1.1.529 with other major SARS-CoV-2 variants. **b**, Key spike mutations found in the viruses isolated in the major lineage of B.1.1.529 are denoted. **c**, Neutralization of D614G and B.1.1.529 pseudoviruses by convalescent patient sera. **d**, Neutralization of D614G and B.1.1.529 pseudoviruses by vaccinee sera. Within the four standard vaccination groups, individuals that were vaccinated without documented infection are denoted as circles and individuals that were both vaccinated and infected are denoted as triangles. Within the boosted group, Moderna vaccinees are denoted as squares and Pfizer vaccinees are denoted as diamonds. **e**, Neutralization of authentic D614G and B.1.1.529 viruses by vaccinee sera. Moderna vaccinees are denoted as squares and Pfizer vaccinees are denoted as diamonds. Data represent one of two independent experiments. For all panels, values above the symbols denote geometric mean titer and the numbers in parentheses denote the number of samples above the limit of detection. *P* values were determined by using a Wilcoxon matched-pairs signed-rank test (two-tailed).

### Serum neutralization of B.1.1.529

We first examined the neutralizing activity of serum collected in the Spring of 2020 from COVID-19 patients, who were presumably infected with the wild-type SARS-CoV-2 (9-120 days post-symptoms) (see Methods and **Extended Data Table 1**). Samples from 10 individuals were tested for neutralization against both D614G (WT) and B.1.1.529 pseudoviruses. While robust titers were observed against D614G, a significant drop (>32-fold) in ID_50_ (50% infectious dose) titers was observed against B.1.1.529, with only 2 samples showing titers above the limit of detection (LOD) (**Fig. 1c and Extended Data Fig. 2a**). We then assessed the neutralizing activity of sera from individuals who received one of the four widely used COVID-19 vaccines: BNT162b2 (Pfizer, 15-213 days post-vaccination), mRNA-1273 (Moderna, 6-177 days post-vaccination), Ad26.COV2.S (Johnson & Johnson, 50-186 days post-vaccination), and ChAdOx1 nCoV-19 (AstraZeneca, 91-159 days post-vaccination) (see Methods and **Extended Data Table 2**). In all cases, a substantial loss in neutralizing potency was observed against B.1.1.529 (**Fig 1d and Extended Data Fig. 2b-f**)For the two mRNA-based vaccines, BNT162b2 and mRNA-1273, a >21-fold and >8.6-fold decrease in ID_50_ was seen, respectively. We note that, for these two groups, we specifically chose samples with high titers such that the fold-change in titer could be better quantified, so the difference in the number of samples having titers above the LOD (6/13 for BNT162b2 versus 11/12 for mRNA-1273) may be favorably biased. Within the Ad26.COV2.S and ChAdOx1 nCOV-19 groups, all samples were below the LOD against B.1.1.529, except for two Ad26.COV2.S samples from patients with a previous history of SARS-CoV-2 infection (**Fig. 1d**). Collectively, these results suggest that individuals who were previously infected or fully vaccinated remain at risk for B.1.1.529 infection.

**Fig. 2.**
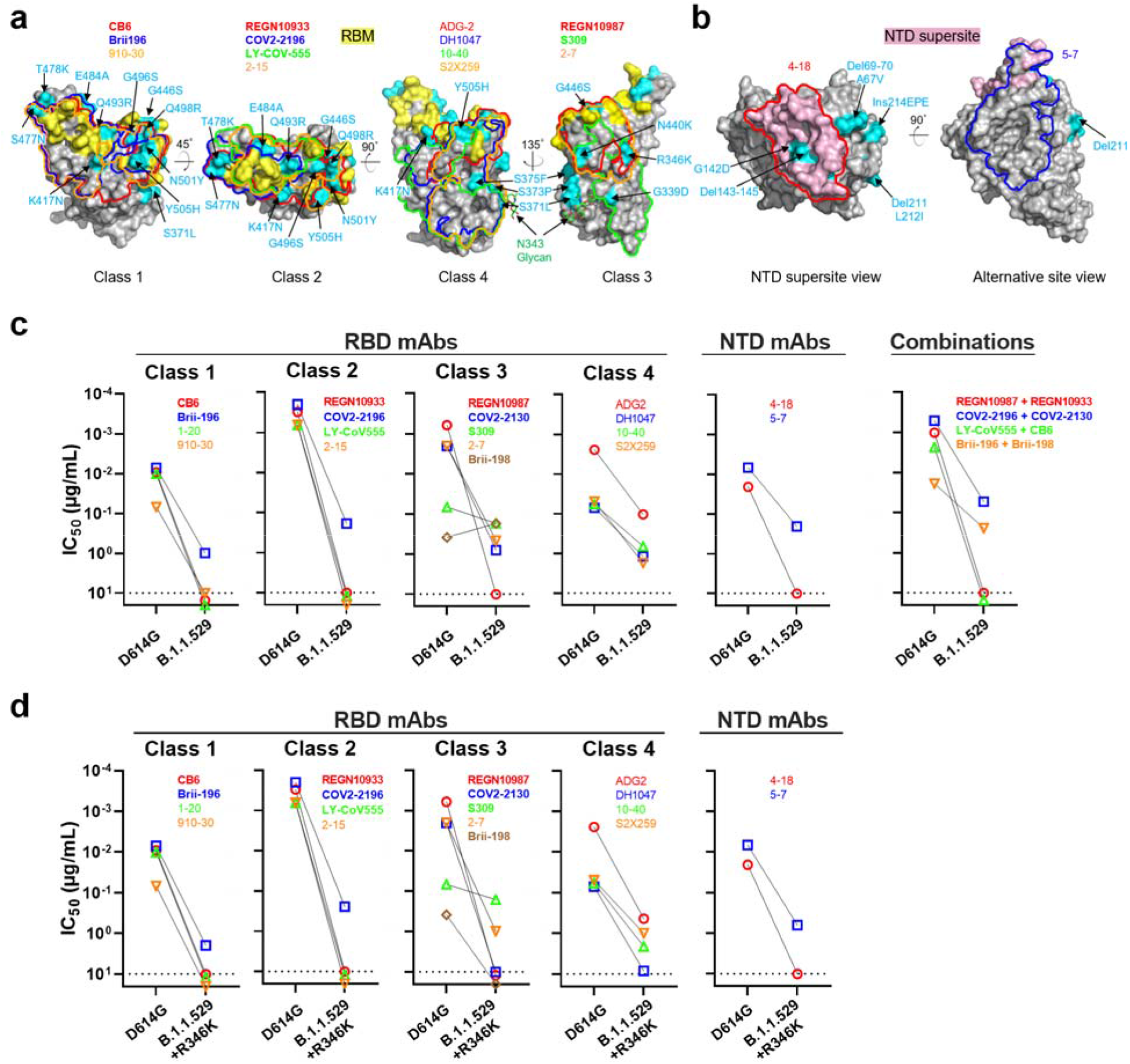
Resistance of B.1.1.529 to neutralization by monoclonal antibodies. **a**, Footprints of RBD-directed antibodies, with mutations within B.1.1.529 highlighted in cyan. Approved or authorized antibodies are bolded. The receptor binding motif (RBM) residues are highlighted in yellow. **b**, Footprints of NTD-directed antibodies, with mutations within B.1.1.529 highlighted in cyan. The NTD supersite residues are highlighted in light pink. **c**, Neutralization of D614G and B.1.1.529 pseudoviruses by RBD-directed and NTD-directed mAbs. **d**, Neutralization D614G and B.1.1.529+R346K pseudoviruses by RBD-directed and NTD-directed mAbs. Data represent one of two independent experiments.

Booster shots are now routinely administered in many countries 6 months after full vaccination. Therefore, we also examined the serum neutralizing activity of individuals who had received three homologous mRNA vaccinations (13 with BNT162b2 and 2 with mRNA-1273, 14-90 days post-vaccination). Every sample showed lower activity in neutralizing B.1.1.529, with a mean drop of 6.5-fold compared to WT (**Fig. 1d**). Although all samples had titers above the LOD, the substantial loss in activity may still pose a risk for B.1.1.529 infection despite the booster vaccination.

We then confirmed the above findings by testing a subset of the BNT162b2 and mRNA-1273 vaccinee serum samples using authentic SARS-CoV-2 isolates: wild type and B.1.1.529. Again, a substantial decrease in neutralization of B.1.1.529 was observed, with mean drops of >6.0-fold and >4.1-fold for the fully vaccinated group and the boosted group, respectively (**Fig. 1e**).

### Antibody neutralization of B.1.1.529

To understand the types of antibodies in serum that lost neutralizing activity against B.1.1.529, we assessed the neutralization profile of 19 well-characterized monoclonal antibodies (mAbs) to the spike protein, including 17 directed to RBD and 2 directed to the N-terminal domain (NTD). We included mAbs that have been authorized or approved for clinical use, either individually or in combination: REGN10987 (imdevimab)^12^, REGN10933 (casirivimab)^12^, COV2-2196 (tixagevimab)^13^, COV2-2130 (cilgavimab)^13^, LY-CoV555 (bamlanivimab)^14^, CB6 (etesevimab)^15^, Brii-196 (amubarvimab)^16^, Brii-198 (romlusevimab)^16^, and S309 (sotrovimab)^17^.

We also included other mAbs of interest: 910-30^18^, ADG-2^19^, DH1047^20^, S2X259^21^, and our antibodies 1-20, 2-15, 2-7, 4-18, 5-7, and 10-40^22-24^. The footprints of mAbs with structures available were drawn in relation to the mutations found in B.1.1.529 RBD (**Fig. 2a**) and NTD (**Fig. 2b**). The risk to each of the 4 classes^25^ of RBD mAbs, as well as to the NTD mAbs, was immediately apparent. Indeed, neutralization studies on B.1.1.529 pseudovirus showed that 17 of the 19 mAbs tested lost neutralizing activity completely or partially (**Fig. 2c and Extended Data Fig. 3**). The potency of class 1 and class 2 RBD mAbs all dropped by >100-fold, as did the more potent mAbs in RBD class 3 (REGN10987, COV2-2130, and 2-7). The activities of S309 and Brii-198 were spared. All mAbs in RBD class 4 lost neutralization potency against B.1.1.529 by at least 10-fold, as did mAb directed to the antigenic supersite^26^ (4-18) or the alternate site^23^ (5-7) on NTD. Strikingly, all four combination mAb drugs in clinical use lost substantial activity against B.1.1.529, likely abolishing or impairing their efficacy in patients.

Approximately 10% of the B.1.1.529 viruses in GISAID^1^ (Global Initiative on Sharing All Influenza Data) also contain an additional RBD mutation, R346K, which is the defining mutation for the Mu (B.1.621) variant^27^. We therefore constructed another pseudovirus (B.1.1.529+R346K) containing this mutation for additional testing using the same panel of mAbs (**Fig. 2d**). The overall findings resembled those already shown in **Fig. 2c**, with the exception that the neutralizing activity of Brii-198 was abolished. In fact, nearly the entire panel of antibodies was essentially rendered inactive against this minor form of the Omicron variant.

The fold changes in IC_50_ of the mAbs against B.1.1.529 and B.1.1.529+R346K relative to D614G are summarized in the first two rows of **Fig. 3a**. The remarkable loss of activity observed for all classes of mAbs against B.1.1.529 suggest that perhaps the same is occurring in the serum of convalescent patients and vaccinated individuals.

**Fig. 3.**
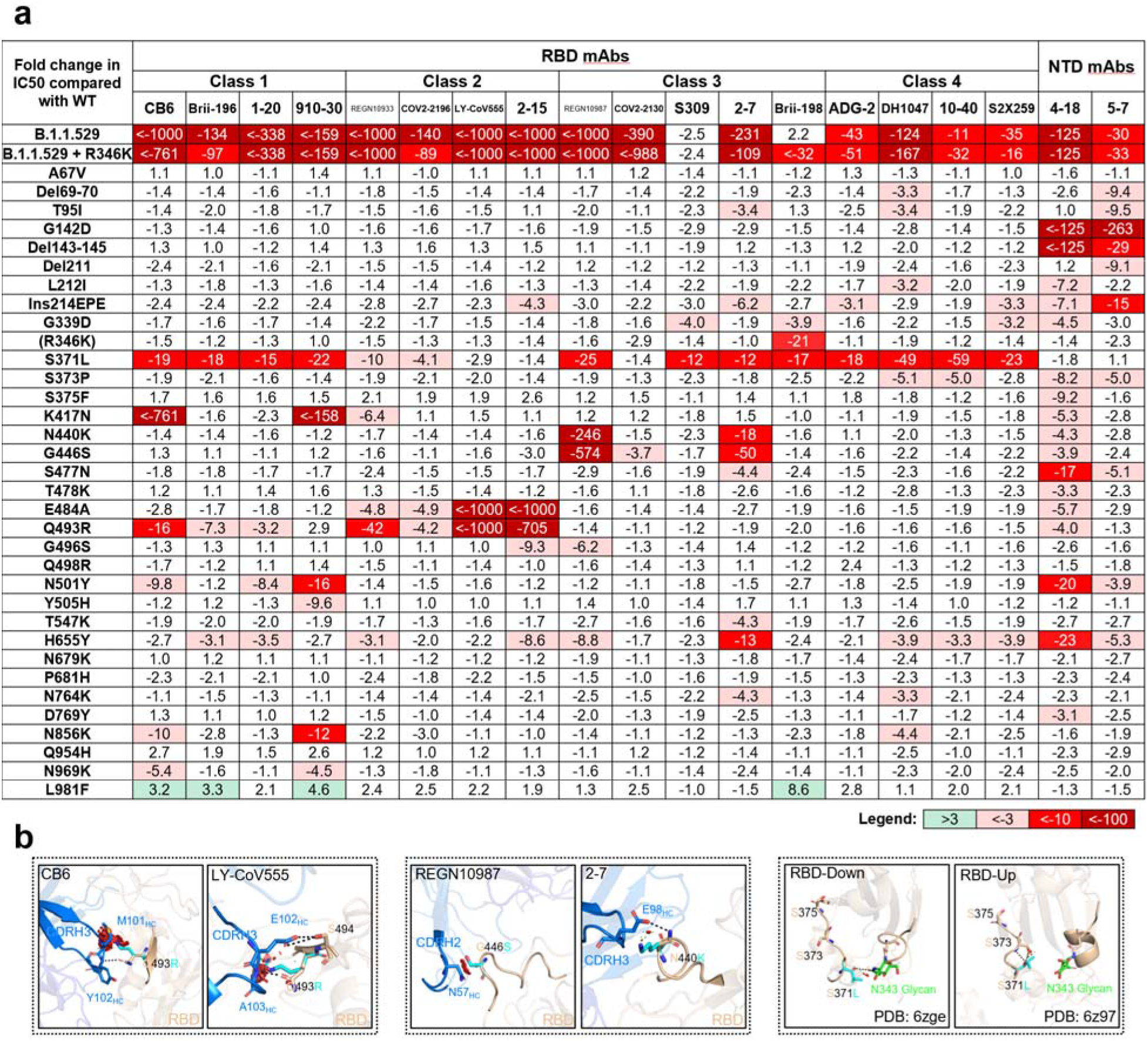
Impact of individual mutations within B.1.1.529 against monoclonal antibodies. **a**, Neutralization of pseudoviruses harboring single mutations found within B.1.1.529 by a panel of 19 monoclonal antibodies. Fold change relative to neutralization of D614G is denoted, with resistance colored red and sensitization colored green. **b**, Modeling of critical mutations in B.1.1.529 that affect antibody neutralization.

### Mutations conferring antibody resistance

To understand the specific B.1.1.529 mutations that confer antibody resistance, we next tested individually the same panel of 19 mAbs against pseudoviruses for each of the 34 mutations (excluding D614G) found in B.1.1.529 or B.1.1.529+R346K. Our findings not only confirmed the role of known mutations at spike residues 142-145, 417, 484, and 501 in conferring resistance to NTD or RBD (class 1 or class 2) antibodies^4^ but also revealed several mutations that were previously not known to have functional importance to neutralization (**Fig. 3a and Extended Data Fig. 4**). Q493R, previously shown to affect binding of CB6 and LY-CoV555^28^ as well as polyclonal sera^29^, mediated resistance to CB6 (class 1) as well as to LY-CoV555 and 2-15 (class 2), findings that could be explained by the abolishment of hydrogen bonds due to the long side chain of arginine and induced steric clashes with CDRH3 in these antibodies (**Fig. 3b, left panels**). Both N440K and G446S mediated resistance to REGN10987 and 2-7 (class 3), observations that could also be explained by steric hindrance (**Fig. 3b, middle panels**). The most striking and perhaps unexpected finding was that S371L broadly affected neutralization by mAbs in all 4 RBD classes (**Fig. 3a and Extended Data Fig. 4**). While the precise mechanism of this resistance is unknown, in silico modeling suggested two possibilities (**Fig. 3b, right panels**). First, in the RBD-down state, mutating Ser to Leu results in an interference with the N343 glycan, thereby possibly altering its conformation and affecting class 3 antibodies that typically bind this region. Second, in the RBD-up state, S371L may alter the local conformation of the loop consisting of S371-S373-S375, thereby affecting the binding of class 4 antibodies that generally target a portion of this loop^24^. It is not clear how class 1 and class 2 RBD mAbs are affected by this mutation.

### Evolution of SARS-CoV-2 to antibodies

To gain insight into the antibody resistance of B.1.1.529 relative to previous SARS-CoV-2 variants, we evaluated the neutralizing activity of the same panel of neutralizing mAbs against pseudoviruses for B.1.1.7^8^, B.1.526^30^, B.1.429^31^, B.1.617.2^9^, P.1^32^, and B.1.351^33^. It is evident from these results (**Fig. 4** and **Extended Data Fig. 5**) that previous variants developed resistance only to NTD antibodies and class 1 and class 2 RBD antibodies. Here B.1.1.529, with or without R346K, has made a big mutational leap by becoming not only nearly completely resistant to class 1 and class 2 RBD antibodies, but also substantial resistance to both class 3 and class 4 RBD antibodies. B.1.1.529 is now the most complete “escapee” from neutralization by currently available antibodies.

**Fig. 4.**
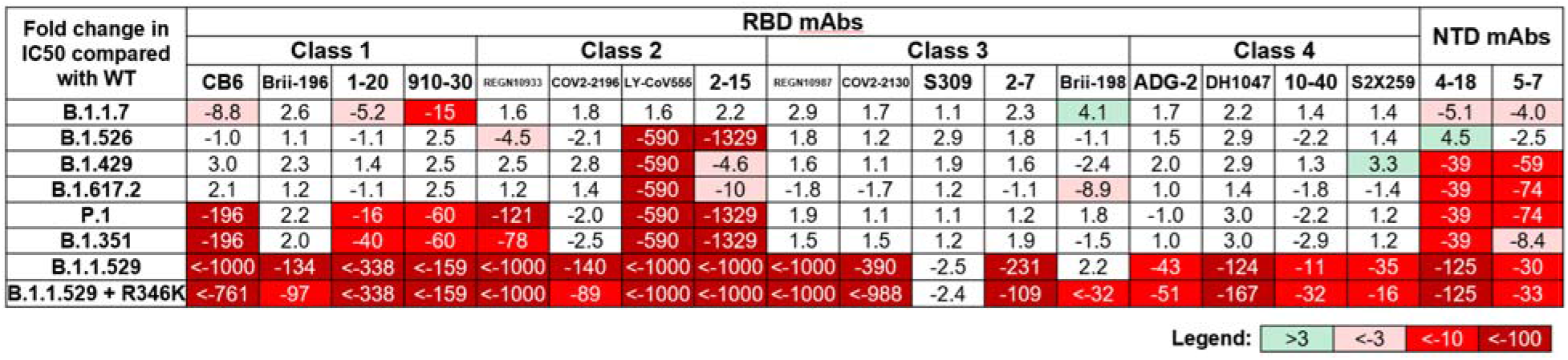
Evolution of antibody resistance across SARS-CoV-2 variants. Neutralization of SARS-CoV-2 variant pseudoviruses by a panel of 19 monoclonal antibodies. Fold change relative to neutralization of D614G is denoted.

## Discussion

The Omicron variant struck fear almost as soon as it was detected to be spreading in South Africa. That this new variant would transmit more readily has come true in the ensuing weeks^2^. The extensive mutations found in its spike protein raised concerns that the efficacy of current COVID-19 vaccines and antibody therapies might be compromised. Indeed, in this study, sera from convalescent patients (**Fig. 1c**) and vaccinees (**Figs. 1d and 1e**) showed markedly reduced neutralizing activity against B.1.1.529. Other studies have found similar losses^34-38^. These findings are in line with emerging clinical data on the Omicron variant demonstrating higher rates of reinfection^11^ and vaccine breakthroughs. In fact, recent reports showed that the efficacy of two doses of BNT162b2 vaccine has dropped from over 90% against the original SARS-CoV-2 strain to approximately 40% and 33% against B.1.1.529 in the United Kingdom^39^ and South Africa^40^, respectively. Even a third booster shot may not adequately protect against Omicron infection^39,41^, although the protection against disease still makes it advisable to administer booster vaccinations. Vaccines that elicited lower neutralizing titers^35,42^ are expected to fare worse against B.1.1.529.

The nature of the loss in serum neutralizing activity against B.1.1.529 could be discerned from our findings on a panel of mAbs directed to the viral spike. The neutralizing activities of all four major classes of RBD mAbs and two distinct classes of NTD mAbs are either abolished or impaired (**Figs. 2c and 2d**). In addition to previously identified mutations that confer antibody resistance^4^, we have uncovered four new spike mutations with functional consequences. Q493R confers resistance to some class 1 and class 2 RBD mAbs; N440K and G446S confer resistance to some class 3 RBD mAbs; and S371L confers global resistance to many RBD mAbs via mechanisms that are not yet apparent. While performing these mAb studies, we also observed that nearly all the currently authorized or approved mAb drugs are rendered weak or inactive by B.1.1.529 (**Figs. 2c and 3a**). In fact, the Omicron variant that contains R346K almost flattens the antibody therapy landscape for COVID-19 (**Fig. 2d and 3a**).

The scientific community has chased after SARS-CoV-2 variants for a year. As more and more of them appeared, our interventions directed to the spike became increasingly ineffective. The Omicron variant has now put an exclamation mark on this point. It is not too far-fetched to think that this SARS-CoV-2 is now only a mutation or two away from being pan-resistant to current antibodies, either monoclonal or polyclonal. We must devise strategies that anticipate the evolutional direction of the virus and develop agents that target better conserved viral elements.

## Figure Legends

**Extended Data Fig. 1**. Mutations within B.1.1.529 denoted on the full SARS-CoV-2 spike trimer (PDB: 6zge).

**Extended Data Fig. 2**. Individual neutralization curves for pseudovirus neutralization assays by serum. Neutralization by **a**, convalescent sera. **b**, Pfizer (BNT162b2) vaccinee sera. **c**, Moderna (mRNA-1273) vaccinee sera. **d**, J&J (Ad26.COV2.S) vaccinee sera. **e**, AstraZeneca (ChAdOx1 nCoV-19) vaccinee sera. **f**, boosted (three homologous BNT162b2 or mRNA-1273 vaccinations) vaccinee sera. Error bars denote mean ± standard error of the mean (SEM) for three technical replicates.

**Extended Data Fig. 3**. Individual neutralization curves for pseudovirus neutralization assays by monoclonal antibodies. Error bars denote mean ± standard error of the mean (SEM) for three technical replicates.

**Extended Data Fig. 4**. Individual neutralization curves for pseudovirus neutralization assays by monoclonal antibodies against individual SARS-CoV-2 mutations. Error bars denote mean ± standard error of the mean (SEM) for three technical replicates.

**Extended Data Fig. 5**. Individual neutralization curves for pseudovirus neutralization assays by monoclonal antibodies against SARS-CoV-2 variants. Error bars denote mean ± standard error of the mean (SEM) for three technical replicates.

**Extended Data Table 1**. Demographics and vaccination information for serum samples from convalescent patients used in this study.

**Extended Data Table 2**. Demographics and vaccination information for serum samples from vaccinated individuals used in this study.

**Extended Data Table 3**. Oligos used to construct spike expression plasmids.

## Methods

### Data reporting

No statistical methods were used to predetermine sample size. The experiments were not randomized and the investigators were not blinded to allocation during experiments and outcome assessment.

### Serum samples

Convalescent plasma samples were obtained from patients with documented SARS-CoV-2 infection. These samples were collected at the beginning of the pandemic in early 2020 at Columbia University Irving Medical Center, and therefore are assumed to be infection by the wild-type strain of SARS-CoV-2^4^. Sera from individuals who received two or three doses of mRNA-1273 or BNT162b2 vaccine were collected at Columbia University Irving Medical Center at least two weeks after the final dose. Sera from individuals who received one dose of Ad26.COV2.S or two doses of ChAdOx1 nCov-19 were obtained from BEI Resources. Some individuals were also infected by SARS-CoV-2 in addition to the vaccinations they received. Note that, whenever possible, we specifically chose samples with high titers against the wild-type strain of SARS-CoV-2 such that the loss in activity against B.1.1.529 could be better quantified, and therefore the titers observed here should be considered in that context. All collections were conducted under protocols reviewed and approved by the Institutional Review Board of Columbia University. All participants provided written informed consent. Additional information for the convalescent samples can be found in **Extended Data Table 1** and for vaccinee samples can be found in **Extended Data Table 2**.

### Monoclonal antibodies

Antibodies were expressed as previously described^22^, by synthesis of heavy chain variable (VH) and light chain variable (VL) genes (GenScript), transfection of Expi293 cells (Thermo Fisher), and affinity purification from the supernatant by rProtein A Sepharose (GE). REGN10987, REGN10933, COV2-2196, and COV2-2130 were provided by Regeneron Pharmaceuticals, Brii-196 and Brii-198 were provided by Brii Biosciences, CB6 was provided by Baoshan Zhang and Peter Kwong (NIH), and 910-30 was provided by Brandon DeKosky (MIT).

### Cell lines

Expi293 cells were obtained from Thermo Fisher (Catalog #A14527), Vero E6 cells were obtained from ATCC (Catalog# CRL-1586), HEK293T cells were obtained from ATCC (Catalog# CRL-3216), and Vero-E6-TMPRSS2 cells were obtained from JCRB (Catalog# JCRB1819). Cells were purchased from authenticated vendors and morphology was confirmed visually prior to use. All cell lines tested mycoplasma negative.

### Variant SARS-CoV-2 spike plasmid construction

An in-house high-throughput template-guide gene synthesis approach was used to generate spike genes with single or full mutations of B.1.1.529. Briefly, 5’-phosphorylated oligos with designed mutations were annealed to the reverse strand of the wild-type spike gene construct and extended by DNA polymerase. Extension products (forward-stranded fragments) were then ligated together by Taq DNA ligase and subsequently amplified by PCR to generate variants of interest. To verify the sequences of variants, next generation sequencing (NGS) libraries were prepared following a low-volume Nextera sequencing protocol^43^ and sequenced on the Illumina Miseq platform (single-end mode with 50 bp R1). Raw reads were processed by Cutadapt v2.1^44^ with default setting to remove adapters and then aligned to reference variants sequences using Bowtie2 v2.3.4^45^ with default setting. Resulting reads alignments were then visualized in Integrative Genomics Viewer^46^ and subjected to manual inspection to verify the fidelity of variants. Sequences of the oligos used in variants generation are provided in **Extended Data Table 3**.

### Pseudovirus production

Pseudoviruses were produced in the vesicular stomatitis virus (VSV) background, in which the native glycoprotein was replaced by that of SARS-CoV-2 and its variants, as previously described^24^. Briefly, HEK293T cells were transfected with a spike expression construct with polyethylenimine (PEI) (1 mg/mL) and cultured overnight at 37 °C under 5% CO_2_, and then infected with VSV-G pseudotyped ΔG-luciferase (G*ΔG-luciferase, Kerafast) one day post-transfection. Following 2 h of infection, cells were washed three times, changed to fresh medium, and then cultured for approximately another 24 h before supernatants were collected, centrifuged, and aliquoted to use in assays.

### Pseudovirus neutralization assay

All viruses were first titrated to normalize the viral input between assays. Heat-inactivated sera or antibodies were first serially diluted in 96 well-plates in triplicate, starting at 1:100 dilution for sera and 10 μg/mL for antibodies. Viruses were then added and the virus-sample mixture was incubated at 37 °C for 1 h. Vero-E6 cells (ATCC) were then added at a density of 3 × 10^4^ cells per well and plates were incubated at 37 °C for approximately 10 h. Luciferase activity was quantified by using the Luciferase Assay System (Promega) according to the manufacturer’s instructions using the software SoftMax Pro 7.0.2 (Molecular Devices, LLC). Neutralization curves and IC_50_ (50% inhibitory concentration) values were derived by fitting a non-linear five-parameter dose-response curve to the data in GraphPad Prism version 9.2.

### Authentic virus isolation and propagation

Authentic B.1.1.529 was isolated from a specimen from the respiratory tract of a COVID-19 patient in Hong Kong by Kwok-Yung Yuen and colleagues at the Department of Microbiology, The University of Hong Kong. Isolation of wild-type SARS-CoV-2 was previously described^47^. Viruses were propagated in Vero-E6-TMPRSS2 cells and sequence confirmed by next-generation sequencing prior to use.

### Authentic virus neutralization assay

To measure neutralization of authentic SARS-CoV-2 viruses, Vero-E6-TMPRSS2 cells were first seeded in 96 well-plates in cell culture media (Dulbecco’s Modified Eagle Medium (DMEM) + 10% fetal bovine serum (FBS) + 1% penicillin/streptomycin) overnight at 37 °C under 5% CO_2_ to establish a monolayer. The following day, sera or antibodies were serially diluted in 96 well-plates in triplicate in DMEM + 2% FBS and then incubated with 0.01 MOI (multiplicity of infection) of wild-type SARS-CoV-2 or B.1.1.529 at 37 °C for 1 h. Sera were diluted from 1:100 dilution and antibodies were diluted from 10 μg/mL. Afterwards, the mixture was overlaid onto cells and further incubated at 37 °C under 5% CO_2_ for approximately 72 h. Cytopathic effects were then visually assessed in all wells and scored as either negative or positive for infection by comparison to control uninfected or infected wells in a blinded manner. Neutralization curves and IC_50_ values were derived by fitting a non-linear five-parameter dose-response curve to the data in GraphPad Prism version 9.2.

### Antibody footprint analysis and RBD mutagenesis analysis

The SARS-CoV-2 spike structure used for displaying epitope footprints and mutations within emerging strains was downloaded from PDB (PDBID: 6ZGE). The structures of antibody-spike complexes were also obtained from PDB (7L5B for 2-15, 6XDG for REGN10933 and REGN10987, 7L2E for 4-18, 7RW2 for 5-7, 7C01 for CB6, 7KMG for LY-COV555, 7CDI for Brii-196, 7KS9 for 910-30, 7LD1 for DH1047, 7RAL for S2X259, 7LSS for 2-7, and 6WPT for S309). Interface residues were identified using PISA^48^ using default parameters. The footprint for each antibody was defined by the boundaries of all epitope residues. The border for each footprint was then optimized by ImageMagick 7.0.10-31 (https://imagemagick.org). PyMOL 2.3.2 was used to perform mutagenesis and to make structural plots (Schrödinger).

## Acknowledgements

We are grateful Regeneron Pharmaceuticals, B. Zhang and P. Kwong (NIAID), and B. Dekosky (MIT) for antibodies. This study was supported by funding from the Gates Foundation, JPB Foundation, Andrew and Peggy Cherng, Samuel Yin, Carol Ludwig, David and Roger Wu, Health@InnoHK, and the National Science Foundation (MCB-2032259).

## Author contributions

D.D.H. conceived this project. L.H.L., S.I., and M.W. conducted pseudovirus neutralization experiments. J.F-W.C., H.C., K.K-H.C., T.T-T.Y., C.Y., K.K-W.T., and H.C. conducted authentic virus neutralization experiments. Y.G. and Z.Z. conducted bioinformatic analyses. L.Y.L. and Y.M.H. constructed the spike expression plasmids. Y.L. managed the project. J.Y. expressed and purified antibodies. M.T.Y. and M.E.S. provided clinical samples. M.S.N. and Y.X.H. contributed to discussions. H.H.W., K-Y.Y., and D.D.H. directed and supervised the project. L.H.L., S.I., and D.D.H. analyzed the results and wrote the manuscript.

## Data availability

Materials used in this study will be made available under an appropriate Materials Transfer Agreement. All the data are provided in the paper. The structures used for analysis in this study are available from PDB under IDs 6ZGE, 7L5B, 6XDG, 7L2E, 7RW2, 7C01, 7KMG, 7CDI, 7KS9, 7LD1, 7RAL, 7LSS, and 6WPT.

