## Supplemental Material for "Striking Antibody Evasion Manifested by the Omicron Variant of SARS-CoV-2"

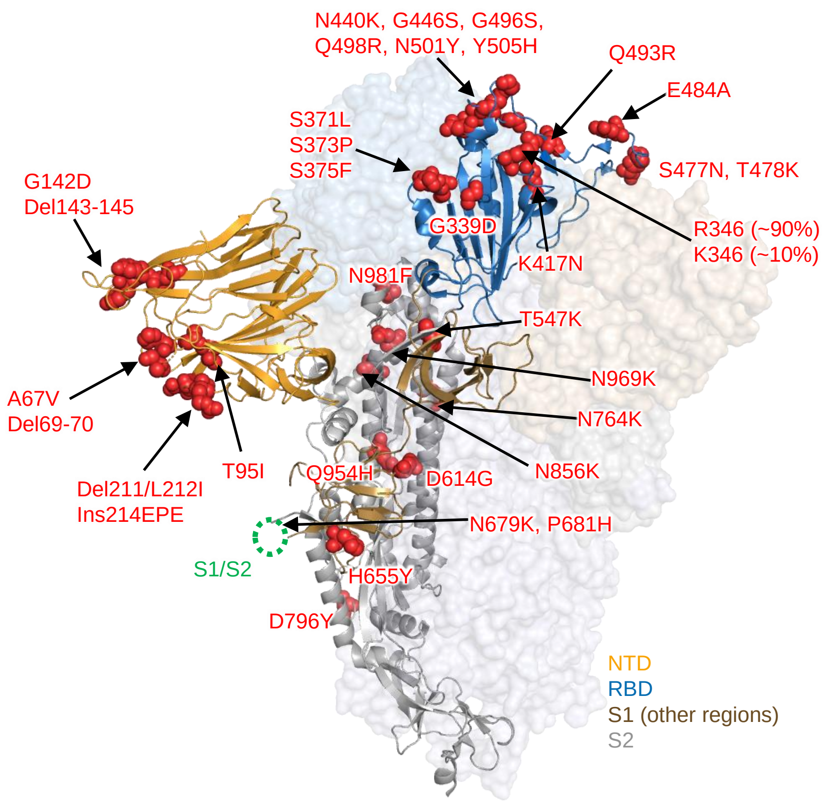


**Extended Data Fig. 1.** Mutations within B.1.1.529 denoted on the full SARS-CoV-2 spike trimer (PDB: 6zge).

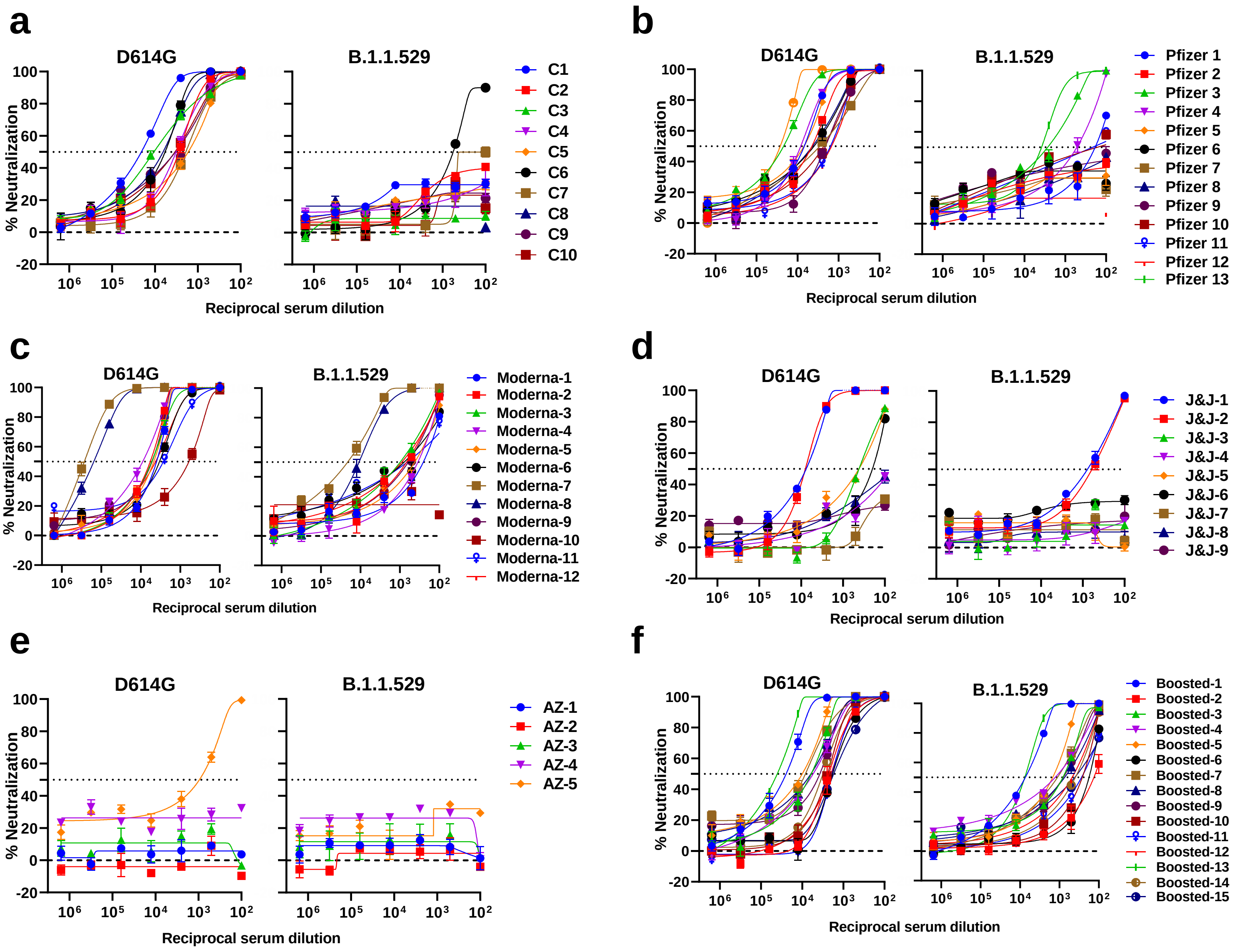


**Extended Data Fig. 2.** Individual neutralization curves for pseudovirus neutralization assays by serum. Neutralization by **a,** convalescent sera. **b,** Pfizer (BNT162b2) vaccinee sera. **c,** Moderna (mRNA-1273) vaccinee sera. **d,** J&J (Ad26.COV2.S) vaccinee sera. **e,** AstraZeneca (ChAdOx1 nCoV-19) vaccinee sera. **f,** boosted (three homologous BNT162b2 or mRNA-1273 vaccinations) vaccinee sera.

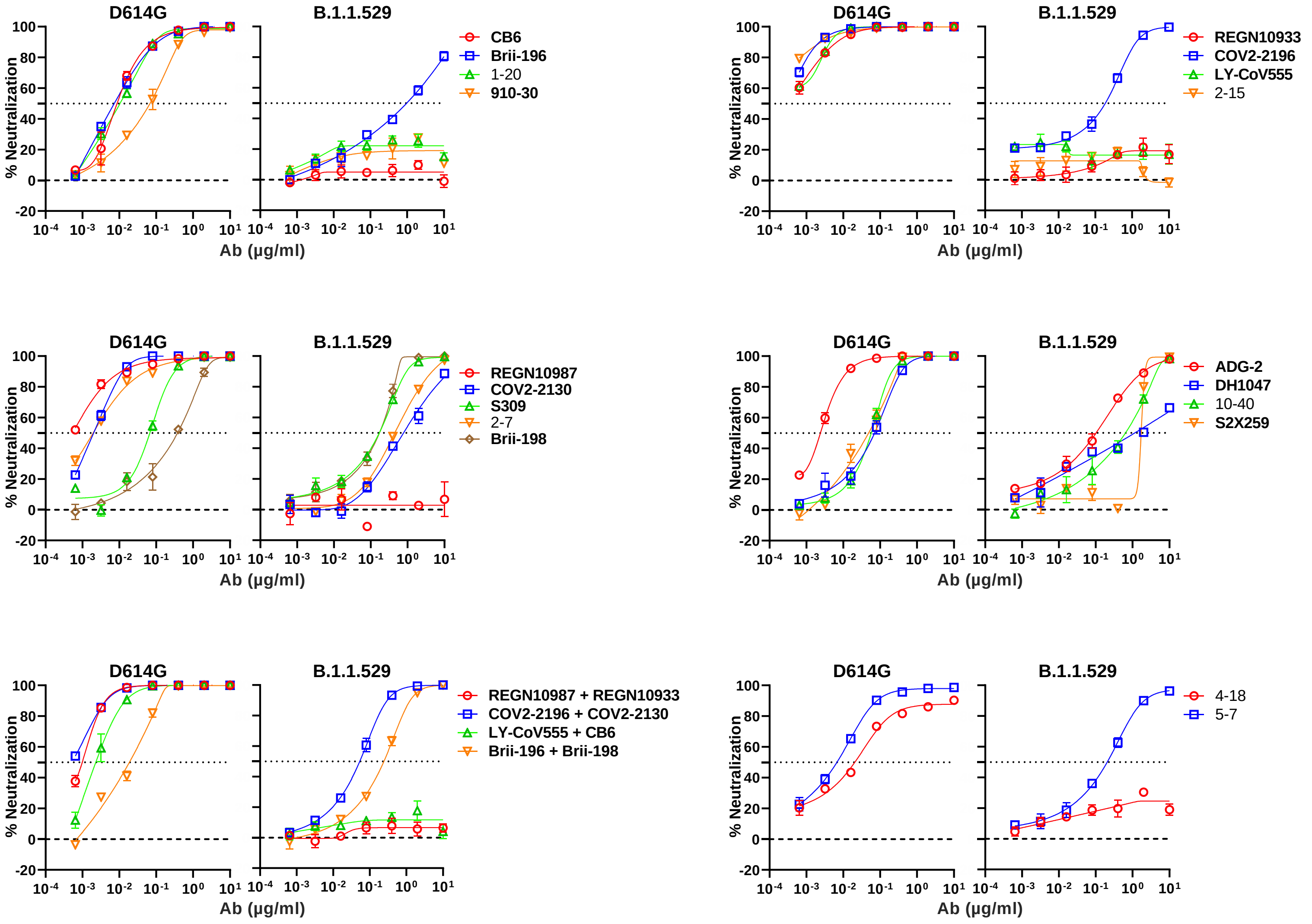


**Extended Data Fig. 3.** Individual neutralization curves for pseudovirus neutralization assays by monoclonal antibodies.

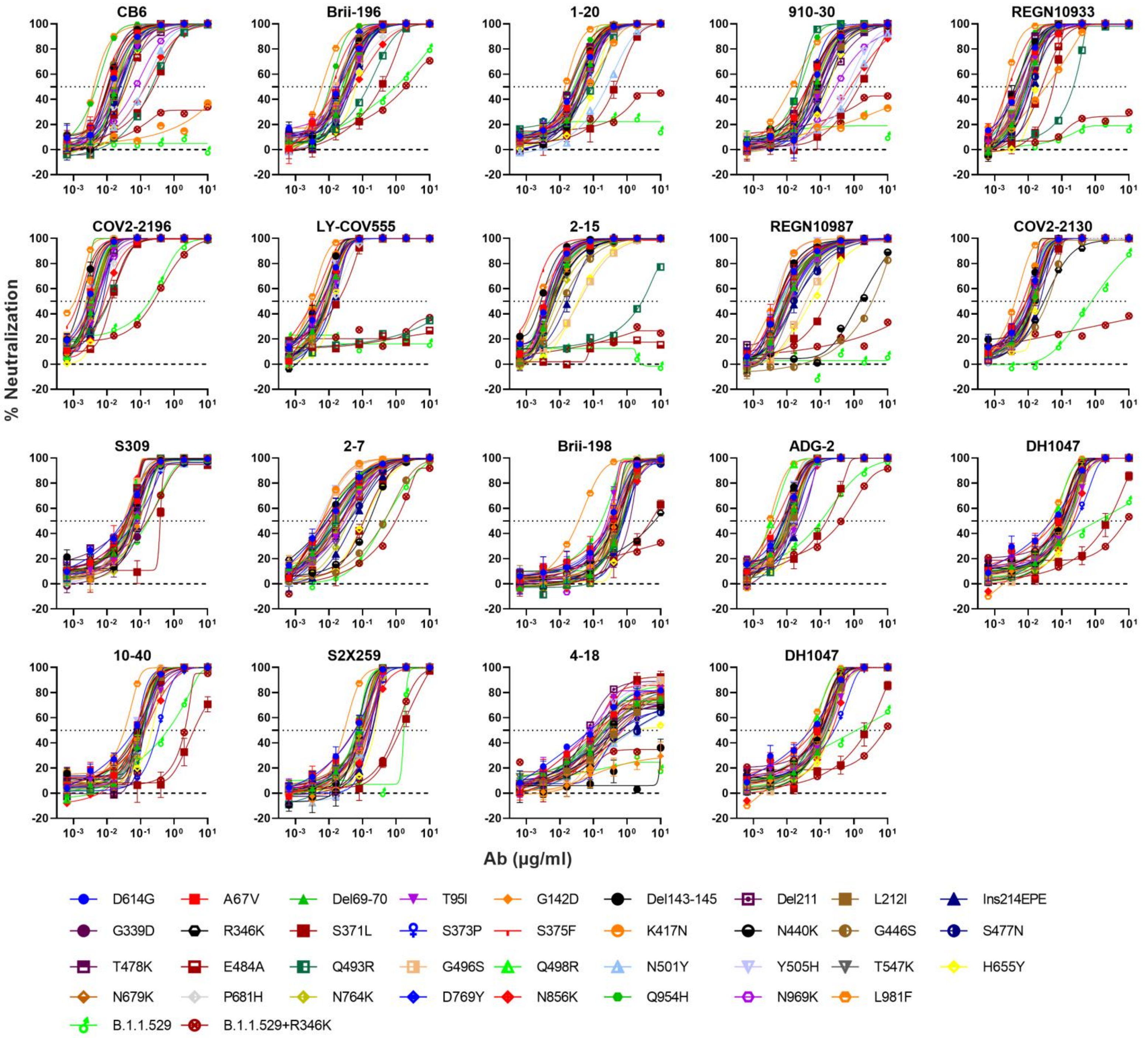


**Extended Data Fig. 4.** Individual neutralization curves for pseudovirus neutralization assays by monoclonal antibodies against individual SARS-CoV-2 mutations.

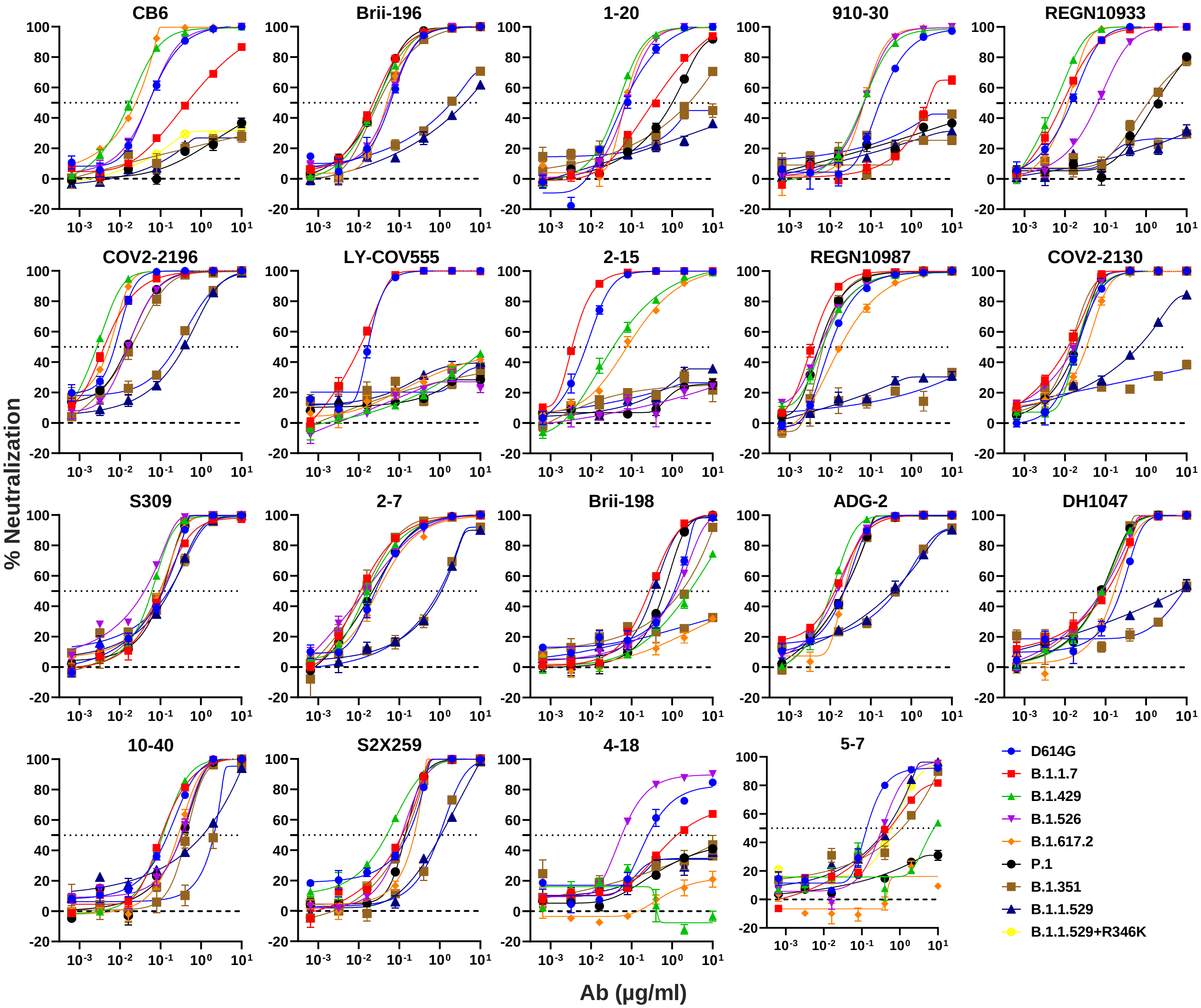


**Extended Data Fig. 5.** Individual neutralization curves for pseudovirus neutralization assays by monoclonal antibodies against SARS-CoV-2 variants.

**Extended Data Table 1.** Demographics and vaccination information for serum samples from convalescent patients used in this study.

| Convalescent Sample | Days post-symptoms | Age | Gender |
|---|---|---|---|
| C1 | 18 | 57 | Female |
| C2 | 25 | 51 | Male |
| C3 | 29 | 71 | Female |
| C4 | 32 | 50 | Male |
| C5 | 35 | 59 | Male |
| C6 | 120 | 56 | Male |
| C7 | 105 | 54 | Female |
| C8 | 77 | 51 | Female |
| C9 | 18 | 79 | Male |
| C10 | 9 | 45 | Male |

**Extended Data Table 2.** Demographics and vaccination information for serum samples from vaccinated individuals used in this study.

| Vaccine Sample | Vaccine type | Days post-vaccination (after last dose) | Documented COVID Infection | Age | Gender |
|---|---|---|---|---|---|
| Moderna vaccinee #1 | mRNA-1273 | 31 | No | 72 | Male |
| Moderna vaccinee #2 | mRNA-1273 | 19 | No | 38 | Female |
| Moderna vaccinee #3 | mRNA-1273 | 6 | No | 42 | Male |
| Moderna vaccinee #4 | mRNA-1273 | 81 | No | 40 | Female |
| Moderna vaccinee #5 | mRNA-1273 | 123 | No | 40 | Female |
| Moderna vaccinee #6 | mRNA-1273 | 177 | No | 40 | Female |
| Moderna vaccinee #7 | mRNA-1273 | 29 | No | 57 | Female |
| Moderna vaccinee #8 | mRNA-1273 | 74 | No | 57 | Female |
| Moderna vaccinee #9 | mRNA-1273 | 32 | No | 66 | Female |
| Moderna vaccinee #10 | mRNA-1273 | 72 | No | 63 | Male |
| Moderna vaccinee #11 | mRNA-1273 | 74 | No | 68 | Female |
| Moderna vaccinee #12 | mRNA-1273 | 58 | No | 46 | Female |
| Pfizer vaccinee #1 | BNT162b2 | 21 | No | 62 | Male |
| Pfizer vaccinee #2 | BNT162b2 | 36 | No | 62 | Male |
| Pfizer vaccinee #3 | BNT162b2 | 26 | No | 38 | Male |
| Pfizer vaccinee #4 | BNT162b2 | 66 | No | 38 | Male |
| Pfizer vaccinee #5 | BNT162b2 | 22 | No | 57 | Female |
| Pfizer vaccinee #6 | BNT162b2 | 61 | No | 57 | Female |
| Pfizer vaccinee #7 | BNT162b2 | 20 | No | 55 | Male |
| Pfizer vaccinee #8 | BNT162b2 | 16 | No | 64 | Female |
| Pfizer vaccinee #9 | BNT162b2 | 32 | No | 68 | Male |
| Pfizer vaccinee #10 | BNT162b2 | 20 | No | 35 | Male |
| Pfizer vaccinee #11 | BNT162b2 | 15 | No | 48 | Female |
| Pfizer vaccinee #12 | BNT162b2 | 21 | No | 45 | Male |
| Pfizer vaccinee #13 | BNT162b2 | 213 | Yes | 66 | Male |
| J&J vaccinee #1 (BEI Cat. #NRH-10818) | Ad26.COV2.S | 55 | Yes | 50 | Female |
| J&J vaccinee #2 (BEI Cat. #NRH-10819) | Ad26.COV2.S | 61 | Yes | 50 | Female |
| J&J vaccinee #3 (BEI Cat. #NRH-10835) | Ad26.COV2.S | 186 | Unknown | 43 | Female |
| J&J vaccinee #4 (BEI Cat. #NRH-10845) | Ad26.COV2.S | 69 | Unknown | 28 | Female |
| J&J vaccinee #5 (BEI Cat. #NRH-10823) | Ad26.COV2.S | 50 | No | 42 | Female |
| J&J vaccinee #6 (BEI Cat. #NRH-10834) | Ad26.COV2.S | 175 | Unknown | 43 | Female |
| J&J vaccinee #7 (BEI Cat. #NRH-10839) | Ad26.COV2.S | 39 | No | 47 | Male |
| J&J vaccinee #8 (BEI Cat. #NRH-10844) | Ad26.COV2.S | 60 | Unknown | 28 | Female |
| J&J vaccinee #9 (BEI Cat. #NRH-10824) | Ad26.COV2.S | 51 | No | 43 | Male |
| AZ vaccinee #1 (BEI Cat. #NRH-10817) | ChAdOx1 nCoV-19 | 158 | Unknown | 73 | Male |
| AZ vaccinee #2 (BEI Cat. #NRH-10814) | ChAdOx1 nCoV-19 | 152 | Unknown | 36 | Female |
| AZ vaccinee #3 (BEI Cat. #NRH-10815) | ChAdOx1 nCoV-19 | 159 | Unknown | 36 | Female |
| AZ vaccinee #4 (BEI Cat. #NRH-10811) | ChAdOx1 nCoV-19 | 142 | Yes | 26 | Female |
| AZ vaccinee #5 (BEI Cat. #NRH-3083) | ChAdOx1 nCoV-19 | 91 | Unknown | 56 | Female |

**Extended Data Table 2 (continued).** Demographics and vaccination information for serum samples from vaccinated individuals used in this study.

| Boosted Sample | Vaccine/boost type | Days post-boost (after last dose) | Documented COVID Infection | Age | Gender |
|---|---|---|---|---|---|
| Boosted sera #1 | mRNA-1273/mRNA-1273 | 28 | No | 66 | Female |
| Boosted sera #2 | BNT162b2/BNT162b2 | 30 | No | 68 | Male |
| Boosted sera #3 | BNT162b2/BNT162b2 | 14 | No | 64 | Female |
| Boosted sera #4 | BNT162b2/BNT162b2 | 34 | No | 55 | Male |
| Boosted sera #5 | BNT162b2/BNT162b2 | 34 | No | 45 | Male |
| Boosted sera #6 | BNT162b2/BNT162b2 | 15 | No | 50 | Female |
| Boosted sera #7 | BNT162b2/BNT162b2 | 15 | No | 48 | Female |
| Boosted sera #8 | BNT162b2/BNT162b2 | 29 | No | 71 | Male |
| Boosted sera #9 | BNT162b2/BNT162b2 | 90 | No | 59 | Male |
| Boosted sera #10 | BNT162b2/BNT162b2 | 33 | No | 45 | Male |
| Boosted sera #11 | BNT162b2/BNT162b2 | 87 | No | 66 | Female |
| Boosted sera #12 | BNT162b2/BNT162b2 | 84 | No | 26 | Male |
| Boosted sera #13 | mRNA-1273/mRNA-1273 | 23 | No | 28 | Female |
| Boosted sera #14 | BNT162b2/BNT162b2 | 14 | No | 78 | Male |
| Boosted sera #15 | BNT162b2/BNT162b2 | 14 | No | 75 | Female |

**Extended Data Table 3.** Oligos used to construct spike expression plasmids.

| Oligo name | Targeted mutations | Oligo sequence |
|---|---|---|
| O_single_mutant1 | A67V | ATGTGACCTGGTTCCATGTGATCCATGTGTCTGGCACCAATGGCACC |
| O_single_mutant2 | Del69-70 | CTGGTTCCATGCCATCTCTGGCACCAATGGCAC |
| O_single_mutant3 | T95I | CTTTGCCAGCATCGAGAAGAGCAACATCATC |
| O_single_mutant4 | Del143-145 | TGTAATGACCCATTCCTGGGACACAAGAACAACAAGTCCTGGATG |
| O_single_mutant5 | G142D | GTAATGACCCATTCCTGGACGTCTACTACCACAAG |
| O_single_mutant6 | Del211 | ACACACACCAATCCTGGTGAGGGACCTG |
| O_single_mutant7 | L212I | CACACCAATCAACATCGTGAGGGACCTGCC |
| O_single_mutant8 | Ins214EPE | ACCAATCAACCTGGTGAGGGAGCCCGAGGACCTGCCACAGGGCTT |
| O_single_mutant9 | G339D | CTGTGTCCATTTGACGAGGTGTTCAATGCCAC |
| O_single_mutant10 | R346K | TGTTCAATGCCACCAAGTTTGCCTCTGTCTATGCCTG |
| O_single_mutant11 | S371F | CTCTGTGCTCTACAACTTTGCCTCCTTCAGCAC |
| O_single_mutant12 | S371L | CTCTGTGCTCTACAACCTGGCCTCCTTCAGCAC |
| O_single_mutant13 | S373P | CTCTACAACTCTGCCCCCTTCAGCACCTTCAAG |
| O_single_mutant14 | S375F | CAACTCTGCCTCCTTCTTCACCTTCAAGTGTTATGG |
| O_single_mutant15 | K417N | CCCCTGGACAAACAGGCAACATTGCTGACTACAACTACAAACTGC |
| O_single_mutant16 | N440K | CCTGGAACAGCAACAAGCTGGACAGCAAGGTG |
| O_single_mutant17 | G446S | GGACAGCAAGGTGAGCGGCAACTACAACTAC |
| O_single_mutant18 | S477N | GATTTACCAGGCTGGCAACACACCATGTAATG |
| O_single_mutant19 | T478K | CAGGCTGGCAGCAAGCCATGTAATGGAGTGGA |
| O_single_mutant20 | E484A | GTAATGGAGTGGCCGGCTTCAACTGTTAC |
| O_single_mutant21 | Q493R | GTTACTTTCCACTCAGATCCTATGGCTTCCAAC |
| O_single_mutant22 | G496S | CACTCCAATCCTATAGCTTCCAACCAACCAATG |
| O_single_mutant23 | Q498R | CAATCCTATGGCTTCAGACCAACCAATGGAGTGGG |
| O_single_mutant24 | N501Y | CTTCCAACCAACCTACGGAGTGGGCTACCAACC |
| O_single_mutant25 | Y505H | AATGGAGTGGGCCACCAACCATACAGG |
| O_single_mutant26 | T547K | CTTCAATGGACTGAAGGGCACAGGAGTGCTGAC |
| O_single_mutant27 | H655Y | CTGATTGGAGCAGAGTACGTGAACAACTCCTATG |
| O_single_mutant28 | N679K | CCAGACCCAGACCAAGAGCCCAAGGAGGGCA |
| O_single_mutant29 | P681H | CCCAGACCAACAGCAGAAGGAGGGCAAGGTCTGTGGC |
| O_single_mutant30 | N764K | GTACCCAACTTAAGAGGGCTCTGACAGGC |
| O_single_mutant31 | D769Y | GACACCTCCAATCAAGTACTTTGGAGGCTTC |
| O_single_mutant32 | N856K | GTGCCCAGAAGTTCAAGGGACTGACAGTGCTG |
| O_single_mutant33 | Q954H | CAAGATGTGGTGAACCACAATGCCCAGGCTCTG |
| O_single_mutant34 | N969K | GCAACTTTCCAGCAAGTTTGGAGCCATCTCCTC |
| O_single_mutant35 | L981F | GTGCTGAATGACATCTTCAGCAGACTGGACAAGGTGGAGG |
| O_multiple_oligo1 | A67V, Del69-70 | TGGTTCCATGTGATCTCTGGCACCAATGG |
| O_multiple_oligo2 | T95I | CTTTGCCAGCATCGAGAAGAGCAAC |
| O_multiple_oligo3 | G142D, Del143-145 | GACCCATTCCTGGACCACAAGAACAACAAGTC |
| O_multiple_oligo4 | L212I, Ins214EPE | CACACACCAATCATCGTGAGGGAGCCCGAGGACCTGCCACAGGGCTTC |
| O_multiple_oligo5 | G339D | TGTGTCCATTTGACGAGGTGTTCAATG |
| O_multiple_oligo6 | S371L, S373P, S375F | TGTGCTCTACAACCTGGCCCCCTTCTTCACCTTCAAGTGTTATG |
| O_multiple_oligo7 | K417N | GGACAAACAGGCAACATTGCTGACTACA |
| O_multiple_oligo8 | N440K, G446S | GCAACAAGCTGGACAGCAAGGTGAGCGGCAACTACAA |
| O_multiple_oligo9 | S477N, T478K, E484A | ACCAGGCTGGCAACAAGCCATGTAATGGAGTGGCCGGCTTCAACTGT |
| O_multiple_oligo10 | Q493R, G496S, Q498R, N501Y, Y505H | TACTTTCCACTCAGATCCTATAGCTTCAGACCAACCTACGGAGTGGGCCACCAACCATACAGGGTGGTGGTGCTGTCCTTTGA |
| O_multiple_oligo11 | T547K | GGACTGAAGGGCACAGGAG |
| O_multiple_oligo12 | D614G | CTCTACCAGGGCGTGAACTGTAC |
| O_multiple_oligo13 | H655Y | TTGGAGCAGAGTACGTGAACAACTC |
| O_multiple_oligo14 | N679K, P681H | CAGACCAAGAGCCACAGGAGGGCAAGG |
| O_multiple_oligo15 | N764K | CCAACTTAAGAGGGCTCTGACAG |
| O_multiple_oligo16 | D796Y | CCTCCAATCAAGTACTTTGGAGGCTTC |
| O_multiple_oligo17 | N856K | CAGAAGTTCAAGGGACTGACAGTGCTG |
| O_multiple_oligo18 | Q954H | GTGGTGAACCACAATGCCCAGGCTC |
| O_multiple_oligo19 | N969K | AACTTTCCAGCAAGTTTGGAGCCATCTCCTC |
| O_multiple_oligo20 | L981F | AATGACATCTTCAGCAGACTGGACAAGGTGGAGGCTGAGGTCCAGATTG |
